# Sustained Ion Imbalances Enhance Ferroptosis

**DOI:** 10.64898/2026.09.30.755323

**Authors:** Ali Yasin Sonay, Elana Apfelbaum, Nermin Mostafa, Jan Grimm

**Author notes:** Corresponding Author: Jan Grimm. Equal contribution first author (Ali Yasin Sonay, Elana Apfelbaum).

## Abstract

Ferroptosis has attracted significant attention for its potential to treat drug resistant tumors, along with its role in neurodegeneration and ischemia. Several ferroptosis targets were identified in the past decade, yet it is also highly affected by cellular metabolism and epithelial or mesenchymal phenotypes which limits the utility of these targets across diverse cell and tissue types. To address this, we utilized chemo-resistant cancer cell lines with different metastatic potential and tissue origins and discovered a common role of cytoplasmic ion changes during ferroptosis. We observed significant changes in intracellular potassium, sodium and chloride levels upon induction of ferroptosis. Therefore, we screened several ion channel inhibitors and activators (i.e. modulators) and identified Kv7.2 and CFTR modulators influence ferroptosis execution despite the fact that these cell lines do not express these proteins. We demonstrated that clinically approved modulators of CFTR and Kv7.2 can inhibit ferroptosis which indicates that these inhibitors could be repurposed as treatments in ferroptosis-associated diseases such as neurodegeneration and kidney diseases independent of ion channel expression. These modulators also influenced potassium and chloride levels in different contexts indicating off-target effects may also lead to changes in ion levels and may serve as a ferroptosis target. Further, we demonstrated that sustained imbalance of sodium, potassium and chloride ions with ferroptosis inducer salinomycin and digoxin, a Na^+^/K^+^ ATPase inhibitor, enhanced ferroptosis. The combination of salinomycin and digoxin successfully enhanced ferroptotic cell death across both cell lines with different metabolic signatures in vitro, offering a novel ferroptosis target. This combination significantly reduced tumor growth in vivo compared with the single agent alone, offering a new therapeutic modality against drug resistant tumors.

## Introduction

Ferroptosis emerged in recent years as a cell death mechanism that can be leveraged as a novel therapeutic modality^1^. During ferroptosis, reactive ferrous iron reacts with hydrogen peroxide via the Fenton reaction to generate hydroxyl radicals. Resulting hydroxyl radicals interact with poly-unsaturated fatty acids in cellular membranes, leading to lipid peroxidation and ultimate cell death by membrane rupture^2^. A cell’s sensitivity to ferroptosis is dependent on cellular antioxidant pathways such as the lipid peroxidation detoxifier GPX4 or cystine uptake symporter Systems Xc-^3^. Inhibition of GPX4 with RSL3 or Systems Xc-with erastin both results in ferroptotic cell death. While ferroptosis is considered a programmed cell death pathway, it is distinct from apoptosis given its caspase independence^2^. This difference made ferroptosis an important novel therapy option against tumors that are highly aggressive and therapy resistant. Moreover, as cells undergo the epithelial to mesenchymal transition (EMT), they develop changes in their lipid metabolism, making them more dependent on GPX4 and susceptible to ferroptosis^4^. As a result, several ferroptosis inducers received significant attention as potential cancer therapies, but their clinical development has been hindered due to their limited efficacy, non-specific toxicity or poor biodistribution and bioavailabilty^5^. These barriers highlight the necessity of discovering novel targets for ferroptosis induction.

One of the challenges in ferroptosis research and its application as therapy is that ferroptosis sensitivity is not universal across cell lines, due to compensatory pathways and metabolic dependencies^6^. Several studies have focused on the enzymatic processes of the ferroptosis pathway and their impact on lipid peroxide species, which has led to the discovery of Protein Kinase C (PKC)^7^, Membrane Bound O-acyl Transferase (MBOAT)^8^ and AMP-Activated Protein Kinase (AMPK)^9^ pathways that are targetable with existing clinically approved drugs. While targeting of these pathways yields significant results, it is often cell type specific since there is a direct interaction between metabolism, cellular energetics and ferroptosis induction^10^. This heterogeneity in ferroptosis machinery often leads to contradictory reports on the involvement of several pathways^11^, and hinders the development of a universal ferroptosis therapy. Beyond serving as a potential cancer therapy, recent studies also showed the role of ferroptosis in physiological and disease processes^12^ such as wound healing^13^, ischemia^14^, neurodegenerative disorders^15^ and finding drug targets for its inhibition also serves as an important healthcare goal.

We aim to find common ferroptosis-associated mechanisms that can be targeted as a more “universal” therapy. We utilized two chemo-resistant, ferroptosis sensitive cell lines (MDA-MB-435 and HT-1080), originating from different tissues to look at fundamental pathways of ferroptotic cell death. As we compared these two cell types, we discovered several common occurrences such as profound changes in cytoplasmic potassium and chloride ion levels during ferroptosis. While previous reports have mentioned calcium sensitive ion channels as potential ferroptosis modulator^16^, we show calcium may influence some cell types but not others. We also identified several ion channel modulators of Kv7.2 and CFTR that can potentially inhibit ferroptosis inhibitors even if the cells do not express these ion channels. Finally, we demonstrate that initiating an ion imbalance with the combination of ferroptosis inducer Salinomycin and clinically approved ion channel modulator Digoxin can enhance ferroptosis and serve as a novel cancer therapy.

## Results

### AMPK and Calcium Enhances Ferroptosis in MDA-MB-435 Cells

Previous studies have emphasized the role of AMPK^9^ and G-Protein Coupled Receptors (GPCRs)^17^ as potential ferroptosis modulators **(Figure 1a)** on a colorectal cancer cell line with limited explorations of upstream and downstream pathway components. To ascertain the effect of distinct cell types and their unique ferroptosis dependencies and compensatory cell protection pathways, we utilized MDA-MB-435 melanoma and HT-1080 fibrosarcoma cancer cell lines. Both of these cell lines are relatively chemo-resistant and sensitive to erastin-induced ferroptosis, with diminished cell death upon treatment with common ferroptosis inhibitors. We first investigated the impact of AMPK, which has been suggested to be involved in ferroptosis, on cell death in these two cell lines. Our results revealed that in MDA-MB-435 cell lines, erastin combined with AMPK activator MK3903 enhanced cell death, whereas AMPK inhibitor Dorsomorphin inhibited erastin induced ferroptosis. Interestingly, HT-1080 cell line did not have a sensitivity towards AMPK modulators **(Figure 1b)**. We further investigated this differing AMPK sensitivity using inhibitors of upstream AMPK pathway components **(Figure 1a).** Our results revealed that inhibition of PKA with H89 or adenylyl cyclase with MDL-12330A led to an increase in erastin-induced cell death in MDA-MB-435 cells, but not in HT-1080 **(Figure 1c).** In contrast, inhibition of other PKA targets such as EPAC did not modulate the ferroptosis induction, indicating this effect is AMPK-dependent **(Supplementary Figure 1a).** Overall, MDA-MB-435 showed increased sensitivity to AMPK pathway modulators compared to HT-1080. This highlights the distinct responses of different cell lines to ferroptosis-associated treatments.

**Figure 1.**
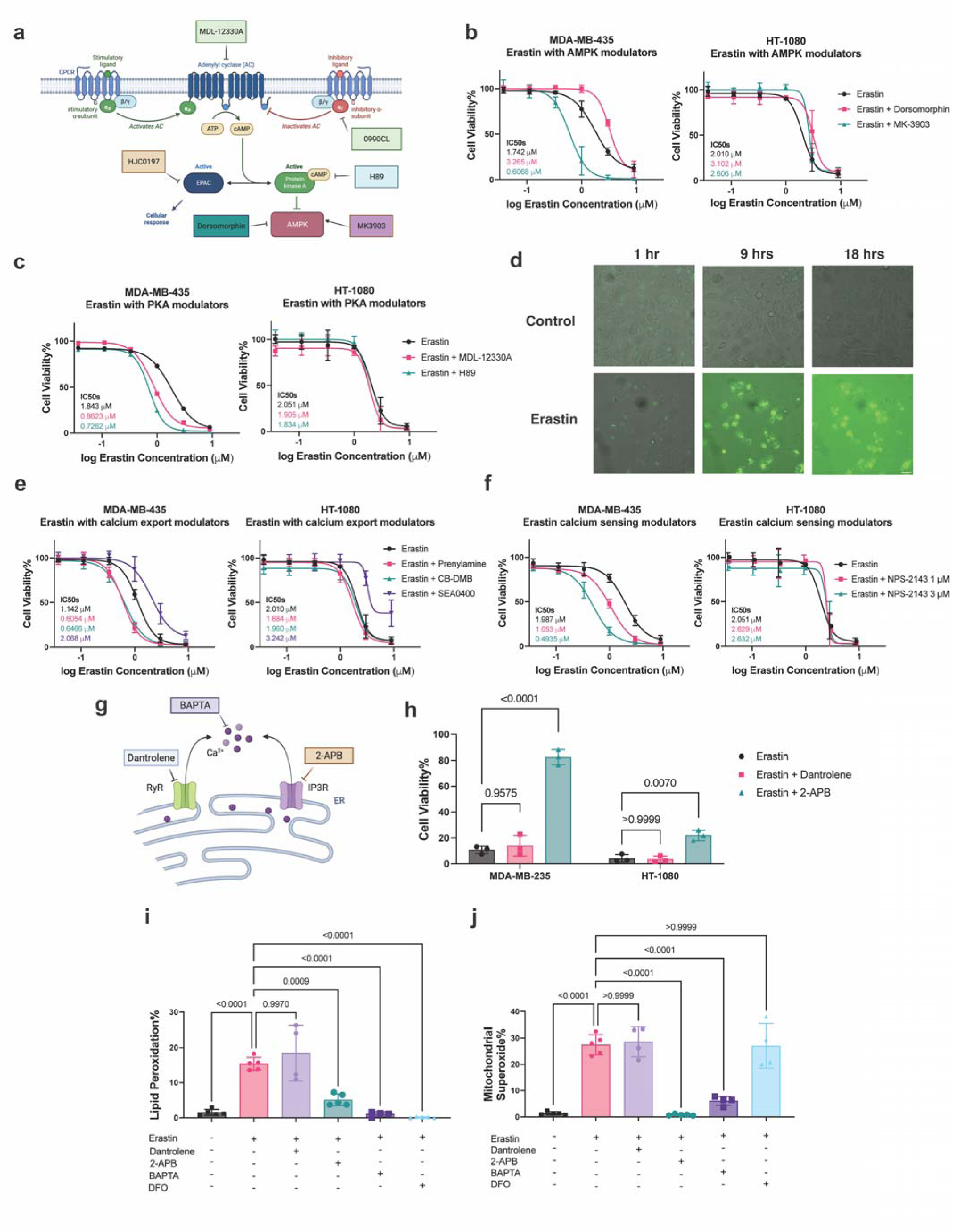
AMPK and Calcium release are important components of ferroptosis but are not part of ferroptosis machinery in all cell lines. **a)** Figure outlining key components of GPCR-PKA-AMPK pathway along with the inhibitors of the key components shown in rectangular boxes, **b)** Erastin induced cell death is enhanced by AMPK activator MK3903 (**5** μ**M**) but decreased by AMPK inhibitor Dorsomorphin (**5** μ**M**) in MDA-MB-435 cell lines(left panel) but not in HT-1080 cell lines(right panel), **c)** Combination of erastin with Adenylyl Cyclase inhibitor MDL-12330A (**5** μ**M**) and PKA inhibitor H89 (**5** μ**M**) lead to increased cell death in MDA-MB-435 cells(left panel), but does not affect erastin induced cell death in HT-1080(right panel). **d)** Live cell imaging of control erastin treated MDA-MB 435 cells with calcium responsive Fluo3-AM dye demonstrate sustained calcium release over time, **e)** In MDA-MB-435 cells, cytoplasmic calcium channel exporters Ca^+2^ ATPase and Na^+^-Ca^+2^ exchanger in forward mode inhibited with Prenylamine (**1** μ**M**) and CB-DMB (**0.25** μ**M**) respectively lead to increased cell death whereas inhibitor of Na^+^-Ca^+2^ exchanger in reverse mode SEA0400 (**10** μ**M**) lead to ferroptosis inhibition(left panel), in HT-1080 cells, inhibition of cytoplasmic calcium channel exporters do not affect erastin induced cell death while whereas inhibitor of Na^+^-Ca^+2^ exchanger in reverse mode SEA0400 prevents ferroptosis(right panel), **f)** Inhibition of Calcium sensing receptor with NPS-2143 (**1** μ**M**) enhances Erastin induced cell death in MDA-MB-435 (left panel), but not in HT-1080 cells (right panel), **g)** Figure showing the RyR and IP_3_R on ER surface acting as calcium channels with inhibitors, **h)** Erastin induced cell death is rescued by IP_3_ receptor inhibitor 2-APB (**10** μ**M**) but not by RyR inhibitor dantrolene (**5** μ**M**) , **i**) Erastin induced lipid peroxidation is inhibited by IP3R inhibitor 2-APB, calcium chelator BAPTA similar to iron chelator DFO, **j)** Erastin incubation leads to increased mitochondrial superoxide levels which are inhibited by IP3R inhibitor and calcium chelator but not DFO. For cytotoxicity assays n=3 biological replicates, for the lipid peroxidation and mitochondrial superoxide assays N=6 pooled from 3 independent experiments. Mean ± s.d. p-values are reported above the lines (Ordinary one-way ANOVA with Tukey’s multiple comparisons). In kill curves concentrations were converted to the log scale and the values were fitted as a nonlinear fit using the log(inhibitor) vs. response function with variable slopes and four parameters.

Calcium, which is implicated in ferroptosis induction of certain cell types^18^, interacts with cAMP-PKA-AMPK axis. We observed a sustained calcium release in MDA-MB-435 cells treated with erastin which lasted several hours **(Figure 1d)**. It is important to note higher background signal observed in 9 and 18 h erastin incubation is due to a loss of membrane structural integrity and release of both calcium and calcium-sensitive dye into the extracellular space. Next, we investigated whether this excess calcium in the cells is removed through other mechanisms such as active calcium export^19^. We combined erastin with inhibitors of the Ca^+2^ ATPase and Na^+^-Ca^+2^ exchangers, Prenylamine and CBDMB, respectively, in forward mode (meaning the inhibitor prevents calcium export and sodium import). Our results revealed that inhibiting calcium export with both inhibitors (i.e. increasing intracellular calcium levels) enhanced ferroptosis in MDA-MB-435 cells, but not in HT-1080 cells. This indicates a distinct role of calcium in ferroptosis where its effects differ between different cell types. In contrast, inhibiting the Na^+^-Ca^+2^ exchanger in reverse mode with SEA0400 (meaning the inhibitor prevents calcium import and sodium export) partially rescued both MDA-MB-435 and HT-1080 cells from ferroptosis **(Figure 1e)**.

After observing the role of calcium export in ferroptosis induction, we investigated the role of membrane-bound GPCRs such as calcium sensing receptor (CaSR) which detect extracellular calcium. We observed that inhibition of CaSR with NPS-2143 led to increased ferroptosis in a dose-dependent manner in MDA-MB-435 cells, but not in HT-1080 cells **(Figure 1f).** We further investigated downstream effects of CaSR inhibition and observed that erastin combined with inhibitors of CaSR synergized with PKA and Adenylyl cyclase inhibitors in MDA-MB-435 cells but not in HT-1080 cells **(Supp. Fig. 1b).** Once again, this highlights the differential role of calcium in ferroptosis between cell lines. To understand the source of the calcium release, we treated cells with different endoplasmic reticulum calcium channel inhibitors **(Figure 1g**). While the Ryanodine Receptor (RyR) inhibitor, Dantrolene, does not affect ferroptotic cell death, IP_3_ receptor inhibitor 2-APB rescues MDA-MB-435 cells from ferroptosis and partially rescues HT-1080 cells **(Figure 1g).** This further supports the hypothesis that ferroptosis in the MDA-MB-435 cell line is more calcium-dependent compared with that of HT-1080. Furthermore, in MDA-MB-435 cells, erastin-induced lipid peroxidation can be inhibited using calcium inhibitors, namely IP_3_ receptor (IP_3_R) inhibitor, 2-APB, and calcium chelator, BAPTA, **(Figure 1i)**. Similarly, erastin treatment has been shown to be associated with increased mitochondrial activity^20^. Our results demonstrate that increased mitochondrial superoxide levels can be inhibited with IP_3_R inhibition and calcium chelation, but, surprisingly, not with iron chelation. This indicates that calcium, not iron, is involved in the ferroptosis-associated mitochondrial activity, while iron is involved in the lipid peroxidation mechanisms **(Figure 1j).** While both ions are crucial for ferroptosis to proceed, this data highlight the distinct roles of iron and calcium in cellular reactive oxygen species production.

### Changes in Cytoplasmic Sodium, Potassium and Chloride Levels during Ferroptosis

Metabolic differences between cell types not only result in distinct calcium dependencies in ferroptosis responses but also lead to variable sensitivities to ferroptosis inducers erastin and RSL3. While both inducers share common features, there are also differences between the molecules and their respective targets which hinder the specificity of ferroptosis-based therapies^1^. To identify universal therapies that are applicable for various cancer types, we investigated common features of ferroptosis that are consistent across different cell lines and ferroptosis inducers. One of the unique features of ferroptosis that sets it apart from apoptotic or necrotic cell death is the membrane blebbing and swelling that eventually leads to a burst and cell death^21^. There is little focus on the causes of this swelling, as it is seen as a natural result of accumulated lipid peroxides and membrane damage, and not a distinct event^22^. After observing the way calcium influences ferroptosis, we investigated whether other ions can be involved in ferroptosis as well, and if this would explain observed differences in the membrane dynamics during ferroptosis. We reasoned that calcium ions in the ER may not be released in every cell type, but other common ions, such as sodium, potassium and chloride, may be more universally affected and cause the unique membrane damage in all cells during ferroptosis.

To this end, we utilized ion-sensing dyes to measure levels of cytoplasmic potassium, sodium, chloride, as well as total cell membrane potential. We observed a decrease of cytoplasmic potassium upon 6 hours after erastin treatment and 2 hours after RSL3 treatment (**Figure 2a),** both of which we denote as “early ferroptosis”. We chose these earlier time points in ferroptosis induction since the loss of membrane integrity at later time points can contribute to the bulk changes in ion levels and would be difficult to differentiate. We also showed an increase in sodium levels in erastin-treated cells, whereas there is only minimal change in RSL3-treated cells **(Figure 2b).** Finally, both RSL3 and erastin treatment led to a decrease in cytoplasmic chloride levels **(Figure 2c).** While the exact mechanisms of these ionic changes are hard to pinpoint, since ions are ubiquitous and crucial in many cellular pathways, rapid shifts in ion concentrations have an overall effect on protein or voltage-dependent ion channel function, which would impact the overall cell homeostasis^22,23^. As expected, we also observed an increase in membrane polarization in cells treated with erastin and RSL3 compared to a positive control digoxin **(Figure 2d)**. It is important to note that while we observe these changes in cytoplasmic ion levels and plasma membrane potential, this does not necessitate a release of these ions to the extracellular space, as some of these ions can be sequestered in other organelles such as the mitochondria or lysosome^23^. In addition, a decrease in membrane polarization can be explained by other mechanisms such as lipid peroxidation and phosphatidylserine exposure because of their negative charges which would normally be facing inner leaflet^22^. However, regardless of the mechanism, these induced membrane changes can have a significant impact on cellular homeostasis, membrane integrity, and ultimately result in cell death.

**Figure 2.**
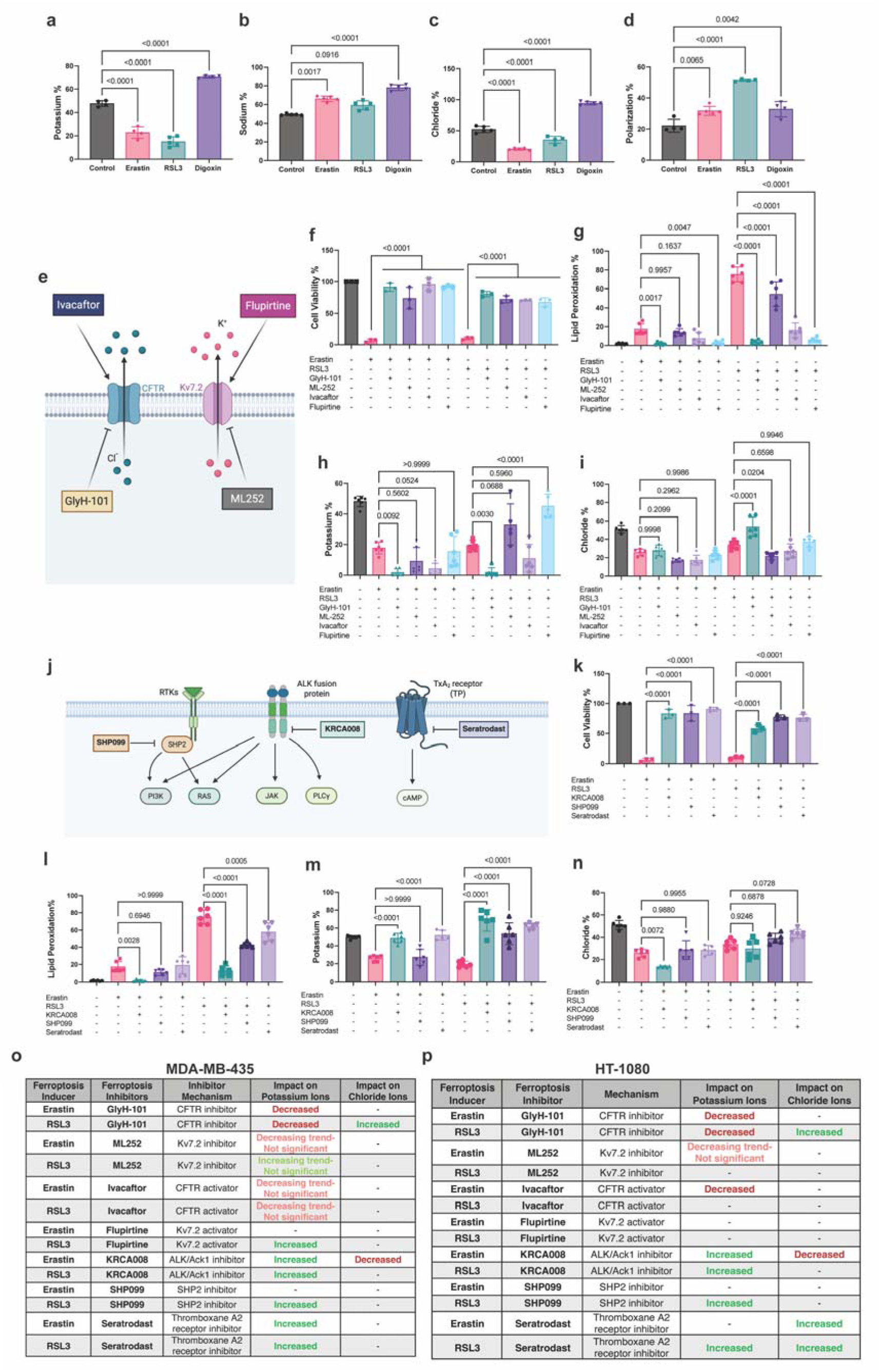
Ferroptosis induction leads to ion imbalances from the early stages and modulation of CFTR and Kv7.2 channels or kinase inhibitors leads to changes in ion levels and ferroptosis inhibition. **a)** Potassium levels in cells treated with erastin, RSL3 and digoxin using IPG-4 AM**, b)** Sodium levels in cells treated with erastin, RSL3 and digoxin using Sodium Green **c)** Chloride levels in cells treated with erastin, RSL3 and digoxin using MQAE dye. **d)** Plasma membrane polarization levels in cells treated with erastin, RSL3 and digoxin using DiBAC_4_. **e)** Figure showing inhibitors and activators of CFTR and Kv7.2 channels, **f)** Cytotoxicity of MDA-MB-435 cells with erastin and RSL3 induced cell death is inhibited by GlyH-101, (2 μM) and activator Ivacaftor(2 μM) , as well as Kv7.2 inhibitor ML252 (2 μM) and activator flupirtine (2 μM). **g)** CFTR and Kv7.2 modulators leads to decreases in the lipid peroxidation levels , **h)** CFTR and Kv7.2 modulators leads to changes in the potassium levels, **i)** CFTR and Kv7.2 modulators leads to changes in the chloride levels in RSL3 induced cell death, but not erastin induced cell death, **j)** Figure showing ALK, SHP2 and Thromboxane A2, their inhibitors and their downstream targets, **k)** Erastin and RSL3 induced cell deaths are inhibited with ALK/Ack1 inhibitor KRCA008 (2 μM), SHP2 inhibitor SHP099 (2 μM), Thromboxane A2 receptor inhibitor Seratrodast (2 μM), **l)** Erastin and RSL3 induced lipid peroxidation are modulated by KRCA008, SHP099, Seratrodast**, m)** Erastin and RSL3 induced potassium level changes are modulated by KRCA008, SHP099, Seratrodast**, n)** Erastin and RSL3 induced chloride level changes are modulated by KRCA008, SHP099, Seratrodast**, o)** Table summarizing the effects of ferroptosis inhibitors on the cytoplasmic potassium and chloride levels in MDA-MB-435 cell line, **p)** Table summarizing the effects of ferroptosis inhibitors on the cytoplasmic potassium and chloride levels in HT-1080 cell line. For cytotoxicity assays n=3 biological replicates, for the lipid peroxidation, potassium, sodium, chloride and membrane polarization levels, N=6 pooled from 3 independent experiments. Mean ± s.d. p-values are reported above the lines (Ordinary one-way ANOVA with Tukey’s multiple comparisons).

After observing the common role of ion imbalance in ferroptosis, we investigated the role of specific ion channels in ferroptotic cell death and screened 7 modulators of 5 ion channels **(Supplementary Figure 2 and 3)**. We discovered that both the CFTR chloride channel^24^ and Kv7.2 voltage gated potassium channel^25^ modulators **(Figure 2e)** rescued the cells from ferroptosis. It is important to highlight neither MDA-MB-435, nor HT-1080 express CFTR and Kv7.2 channels, indicating the observed effect is independent of CFTR or Kv7.2 activity. Yet inhibitors and activators of these channels that we tested rescue from erastin- and RSL3-induced ferroptosis in both MDA-MB-435 **(Figure 2f)** and HT-1080 cells **(Supp. Fig. 4a)**. These results may be explained by potential non-specific effects of these modulators that may impact ferroptosis such as recent reports on ivacaftor’s role in ferroptosis through Nrf2 and GPX4 activation^26^. However, it is also likely that these modulators influence other ion channels and cellular components to influence the ion levels through off-target effects^27,28^. It is important to note that CFTR and Kv7.2 activators ivacaftor^29^ and flupirtine^30^ are both clinically approved and may, therefore, serve as novel ferroptosis inhibitors in future clinical studies. Flupirtine, for instance, is a non-opioid pain killer that is readily prescribed in multiple countries and has been utilized for neuroprotection^31^ and multiple sclerosis^32^, both of which are marked by neuronal cell death components that are associated with ferroptosis.

**Figure 3.**
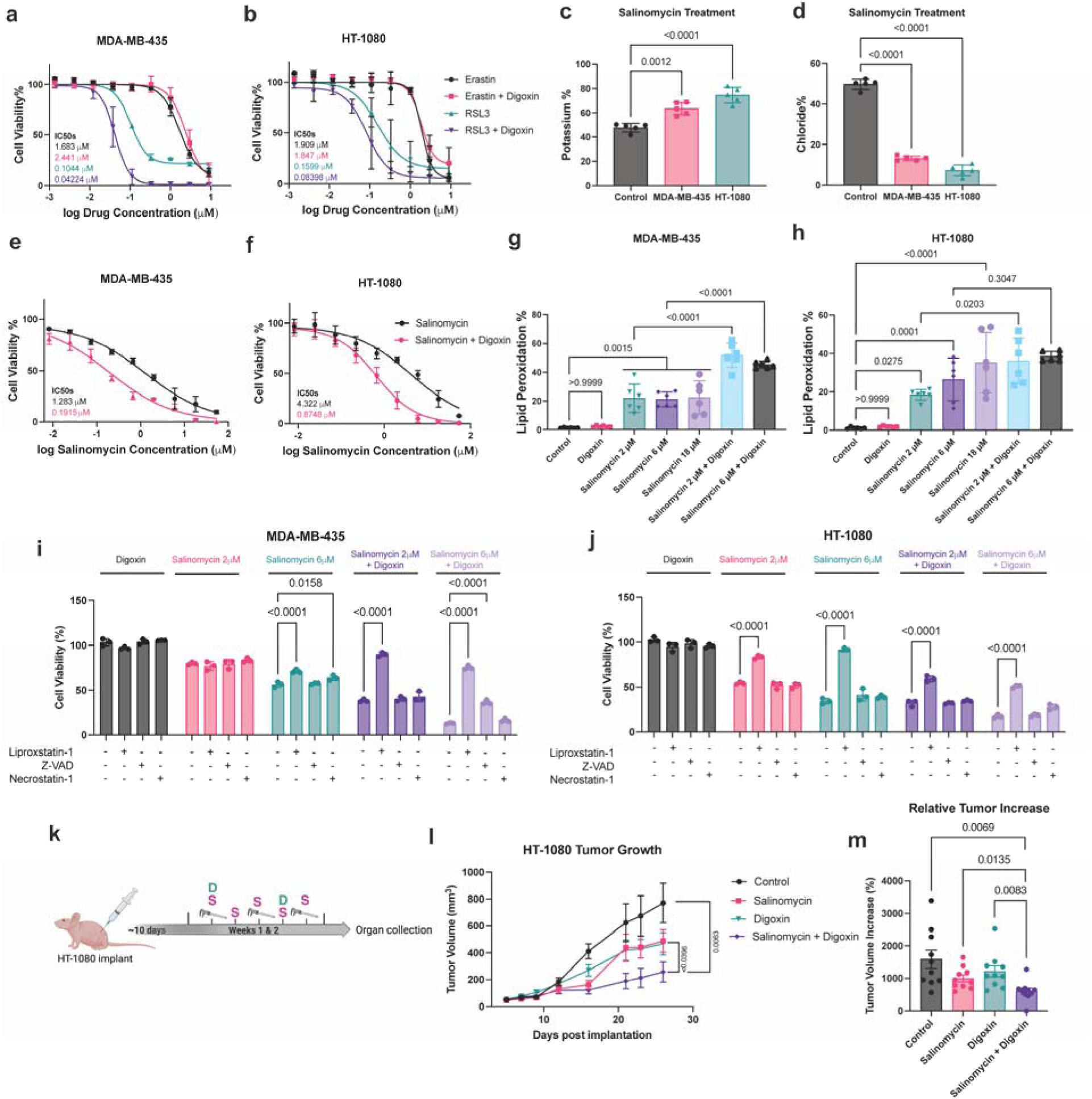
Maintaining Ionic Imbalance with Na^+^ K^+^ ATPase inhibitor Digoxin enhances ferroptosis and its combination with Salinomycin offers a novel treatment strategy. **a)** Combination of digoxin with erastin does not lead to increased cell death but digoxin RSL3 combination leads to enhanced cell death in MDA-MB-435 cells, **b)** Cytotoxicity of erastin and RSL3 alone and combined with digoxin in HT-1080 cells shows RSL3-Digoxin synergy, **c)** Salinomycin induces increases in potassium levels in both MDA-MB-435 and HT-1080 cell lines, **d)** Salinomycin treatment decreases cytoplasmic chloride levels in both MDA-MB-435 and HT-1080 cell lines, **e)** Salinomycin induced cell death either alone or in combination with digoxin in MDA-MB-435 cell lines, **f)** Salinomycin induced cell death either alone or in combination with digoxin in HT-1080 cell lines, **g)** In MDA-MB-435 cells, salinomycin leads to an increase in lipid peroxidation that is not dependent on salinomycin concentration, but is enhanced by salinomycin-digoxin combination, **h)** In HT-1080 cells, salinomycin leads to an increase in lipid peroxidation that is concentration dependent, and enhanced by salinomycin-digoxin combination, **i)** Rescue experiment testing whether cell death induced by digoxin, different concentrations of salinomycin (2 μM and 6 μM) alone or in combination with one another can be rescued by ferroptosis inhibitor liproxstatin, apoptosis inhibitor z-VAD and necroptosis inhibitor necrostatin in MDA-MD-435 melanoma cell lines, **j)** Rescue experiment testing whether cell death induced by digoxin, different concentrations of salinomycin (2 μM and 6 μM) alone or in combination with one another can be rescued by ferroptosis inhibitor liproxstatin, apoptosis inhibitor z-VAD and necroptosis inhibitor necrostatin in HT-1080 fibrosarcoma cell lines, **k)** Experimental workflow of in vivo xenograft therapy denoting the injection frequency of different agents, **l)** Measurements of tumor volumes during the course of the therapy for each treatment group indicating combination therapy showed a more significant decrease in tumor volume compared with control and either therapy alone, **m)** Percent tumor volume increase for different treatment groups show combination therapy prevents tumor growth more significantly compared with all the other therapies.N=10 mice for each group. For cytotoxicity assays n=3 biological replicates, for the lipid peroxidation, potassium and chloride measurements, N=6 pooled from 3 independent experiments. Mean ± s.d. p-values are reported above the lines (Ordinary one-way ANOVA with Tukey’s multiple comparisons). In kill curves concentrations were converted to the log scale and the values were fitted as a nonlinear fit using the log(inhibitor) vs. response function with variable slopes and four parameters.

Given that modulators of CFTR and Kv7.2 protect cells from ferroptotic cell death even in the absence of their target proteins, we tested the effect of these modulators on lipid peroxidation and cytoplasmic ion levels. Our results showed CFTR and Kv7.2 modulators protected both MDA-MB-435 **(Figure 2g)** and HT-1080 cells **(Supplementary Figure 4b)** cell from erastin- and RSL3-induced lipid peroxidation. When we measured the cytoplasmic potassium levels in cells treated with CFTR and Kv7.2 modulators, we consistently observed a decrease in potassium values in response to erastin and RSL3 but to varying degrees. For example, CFTR (chloride) channel inhibitor GlyH-101 leads to a further decrease in cytoplasmic potassium in erastin and RSL3 treated cells, while activation with Ivacaftor leads to a potassium decrease only in erastin-treated HT-1080 cells. On the other hand, Kv7.2 (potassium) channel activator Flupirtine only increases the potassium levels in RSL3 treated MDA-MB-435 cells **(Figure 2h and Supplementary Figure 5a).** While both HT-1080 and MDA-MB-435 cells experience decreased potassium in response to erastin- and RSL3-induced ferroptosis, the roles of Kv7.2 and CFTR modulators in these processes seem to be both cell- and modulator-dependent given their varying off-target effects. In terms of chloride levels, these modulators did not further affect the decreased cytoplasmic chloride from erastin treatment, but CFTR (chloride) channel inhibitor GlyH-101 consistently increased the cytoplasmic chloride levels only in cells treated with RSL3 **(Figure 2i and Supplementary Figure 5b).**

Overall, these results indicate that while some off-target effects are ion channel independent such as ivacaftor’s role in Nrf2 signaling, some may be impacting other ion channels on these cells as observed with potassium flux from chloride channel modulators. While ion dynamics and their role in ferroptosis are a delicate balance and difficult to isolate, these results highlight that ionic changes can be considered a potential target for ferroptosis.

### Effect of ALK, SHP2 and Platelet Activation Inhibitors on Ion Levels

Because ion dynamics play a role in many, if not all, cellular pathways, we investigated other cell signaling pathways that are known to affect these ion levels. To this end, we tested several ferroptosis modulators and observed their effect on ion regulation. Through a small screen, we discovered inhibitors of ALK, TXA2, and SHP2 play important roles in both ferroptosis and its subsequent ionic changes. Previous studies have shown ALK inhibition with crizotinib^30^, SHP2 knockdown^33^ and platelet activation inhibitor Seratrodast^32^ can be utilized to attenuate ferroptotic cell death. Previous work in our lab showed that ferroptosis machinery shares components with platelet activation that might explain this connection^25^. We utilized an ALK/Ack-1 inhibitor (KRCA008), SHP2 inhibitor (SHP099), and TXA2 receptor inhibitor (Seratrodast), **(Figure 2j)** all of which rescued both MDA-MB-435 and HT-1080 cells **(Figure 2k and Supplementary Figure 4c)** from erastin and RSL3 induced ferroptosis. Furthermore, these inhibitors also decreased erastin and RSL3 induced lipid peroxidation, similar to the previous findings **(Figure 2l and Supplementary Figure 4d)**. We further tested the impact of KRCA008, SHP099, Seratrodast on cytoplasmic ion levels, and observed increasing cytoplasmic potassium levels to varying degrees, especially in RSL3-induced ferroptosis in both cell lines **(Figure 2m and Supplementary Figure 5c**). In contrast, they had a milder effect on cytoplasmic chloride concentrations. While KRCA008 enhanced the erastin-mediated chloride decrease in both cell lines, SHP099 and Seratrodast only enhanced RSL3-induced chloride increase in HT-1080 cells **(Figure 2n and Supplementary Figure 5d**). Overall, we highlight the role of ALK, SHP2, and platelet activation pathways in ferroptosis. These protein kinases have downstream effects on ion levels which indicates the close relationship between diverse pathways and ion channels, also their inhibition influences ion homeostasis and rescues the cells from ferroptosis. While the trends are similar, their impact on potassium and chloride levels vary depending on the ferroptosis inducer with which they are combined, as well as cell line. **(Figure 2o and p)**. We observed several common effects of these modulators that are consistent across cell types and ferroptosis inducers, while other modulators had more inconsistent effects.

### Sustained Ion Imbalance as a Ferroptosis Enhancer

Our results demonstrate a complex relationship between ion dynamics and ferroptosis, which is influenced by the ferroptosis inducing agent, cancer type, and metabolic state. We also show that a simple change in one parameter, such as an excess increase or decrease of a specific ion, does not necessarily capture the complex crosstalk between these ion channels and intracellular ion sequestering, which all may lead to different ferroptosis pathways. Ion changes not only influence the membrane dynamics and cell polarity, but also impact the downstream signaling pathways that are essential for cell death^34^. Therefore, we hypothesized that a general sustained ion imbalance, rather than loss of a specific ion type, may be an important parameter in ferroptosis. Our data on ion levels suggest that ion disequilibrium needs to be maintained over time to facilitate accumulation of cellular and membrane damage and initiate ferroptosis, especially since modulation of ion levels in one direction could tip the ion balance the other way and prevent ferroptotic cell death through independent mechanisms.

Based on these findings, we identified drug candidates that could sustain ion imbalance or prevent its recovery could overcome ferroptosis resistance mechanisms, hypothesizing that they may serve as novel ferroptosis inducers or enhancers. To achieve this, we employed the inhibitor of Na^+^/K^+^ ATPase exchanger digoxin, a clinically-approved drug that has been widely used for treatment of arrythmias^35^. While digoxin has been widely used, high doses are extremely toxic, therefore, we investigated digoxin at sub-lethal concentrations. Our results revealed that a combination of RSL3 and digoxin at non-cytotoxic digoxin concentrations led to increased cytoxocity, while combination with erastin did not **(Figure 3a and b).** Based on our cytoplasmic ion level measurements, we knew both erastin and digoxin increased sodium levels individually, whereas RSL3 did not. This could indicate that erastin and digoxin work along the same ion pathway, but digoxin may complement RSL3-induced potassium and chloride release to achieve an enhanced cell death.

Despite these encouraging results, neither erastin, nor RSL3 are clinically approved, which is another large hurdle for moving ferroptosis treatments further. Therefore, we employed a clinically approved antibiotic, salinomycin, which has recently gained interest for selectively targeting breast cancer stem cells^34^ and has been described as a ferroptosis inducer. Although the exact mechanism is under debate, some groups indicate it is a sodium ionophore, whereas others describe it as a loose iron chelator which chelates iron from the lysosome, but then releases it in the cytoplasm so free iron can undergo the Fenton reaction and lipid peroxides^36^. To take full advantage of these ion changes and potential for clinical translation, we investigated salinomycin as a potential pair with digoxin.

We reasoned either of these mechanisms of salinomycin can lead to significant changes in cytoplasmic ion levels. As expected, we saw a significant increase in potassium levels in both cell lines, though more intensely in HT-1080 **(Figure 3c)**. We also observed a decrease in cytoplasmic chloride levels upon salinomycin treatment **(Figure 3d)**.

Finally, we tested whether digoxin can enhance salinomycin induced cell death. Since digoxin cytotoxicity did not increase with increasing concentrations, we could not achieve a dose response curve for digoxin alone precluding the possibility to utilize existing synergy calculations. Therefore we used increasing concentrations of salinomycin either alone or in combination with a fixed sublethal digoxin concentration. We define the dose reduction index to measure the synergy as the ratio of IC_50_ values of salinomycin alone over combination therapy. Our results showed a dose reduction index of 0.149 for MDA-MB-435 cell line and 0.203 for the HT1080 cell line **(Figure 3e and f).** We also investigated whether this combination therapy would lead to increased lipid peroxidation, which is the hallmark of ferroptosis. Salinomycin alone increased lipid peroxidation in both cells, but only in HT-1080 was a dose-dependent increase in lipid peroxidation observed. In MDA-MB-435 Salinomycin-induced lipid peroxidation levels were maintained despite a 3- and 9-fold increase in concentration. Despite these differences, in both cell lines, digoxin alone did not induce lipid peroxidation, while a combination of digoxin and salinomycin significantly enhanced the lipid peroxidation levels **(Figure 3g and h).** This demonstrated the synergistic combination of salinomycin and digoxin, indicating that the sustained ion imbalance can enhance ferroptosis. Furthermore, we tested whether salinomycin alone or digoxin alone would induce ferroptosis, apoptosis and necroptosis by using their inhibitors for each cell line. Our results revealed HT-1080 was more sensitive to ferroptosis treatment and it was readily rescued by ferroptosis inhibitor liproxstatin at different concentrations (2 μM and 6 μM) but not by apoptosis inhibitor z-VAD or necroptosis inhibitor necrostatin **(Figure 3j)**. In contrast, MDA-MB-435 cell lines are rescued by both liproxstatin and necrostatin at higher concentrations of salinomycin (6 μM) **(Figure 3i)**. In both cell lines digoxin showed minimal cytotoxicity. As for the combination therapy of salinomycin and digoxin, we observed an enhanced cell death in both cell lines which were significantly rescued by liproxstatin alone at 2 μM salinomycin combined with fixed digoxin concentration. Interestingly, 6 μM salinomycin with the same digoxin concentration induced cell death that is predominantly rescued by liproxstatin, while other cell death inhibitors also contributed to the rescue, such as z-VAD in MDA-MB-435 cell lines**(Figure 3i)** and necrostatin in HT-1080 cell lines **(Figure 3j)**. It is important to highlight while salinomycin-digoxin combination may not lead to only ferroptotic cell death, such an outcome does not detract from the efficacy of this therapy as tumor tissues often contain cancer cells with various sensitivities against different cell death types. An approach that can target multiple cell death pathways would greatly benefit the outcome of cancer therapy.

After establishing this synergistic combination of ion perturbations and ferroptosis, we investigated this drug combination *in vivo*. Because the HT-1080 model showed the most significant response to the combination *in vitro*, HT-1080 tumor-bearing mice were established. A clinically relevant, but sub-lethal, dose of digoxin (0.5 mg/kg), given biweekly, was combined with daily administration of salinomycin (2.5 mg/kg) **(Figure 3k)**. The combination of salinomycin and digoxin led to significantly decreased tumor growth, compared with the single agents **(Figure 3l and m)**. This therapy resulted in little toxicity, observed with sustained mouse weight and no kidney or liver damage visible in H&E staining (**Supplementary Figure 6)**.

These results highlight the important of ion dynamics in ferroptosis. We show that salinomycin can lead to increased lipid peroxidation and ion level fluctuations *in vitro*, which can be utilized by digoxin. Their combination maintains ion imbalance, leading to enhanced cell death that can be applied in an *in vivo* setting as a potential cancer treatment.

## Discussion

In this study, we highlight how ferroptosis can be influenced by metabolic changes or cell characteristics, and the need for novel therapeutic targets that are consistent across different cell types. Our results revealed cellular ion levels are an important aspect of ferroptosis that has been largely ignored thus far. While calcium changes during the course of ferroptosis are apparently cell type dependent, our results showed consistent changes in cytoplasmic potassium and chloride levels across different cell lines. Our initial hypothesis was that the loss of membrane structural integrity causes these ion level differences while we also explored whether ion changes alter membrane polarization and caused blebbing. Utilizing ferroptosis inhibitors did not rescue these changes, which indicated that ion levels are independent from lipid peroxidation induced damage. To ascertain which ion channels might be implicated in ferroptosis, we utilized a small screen of ion channel modulators and observed little to no change from most of the calcium, potassium, chloride and sodium modulators. Interestingly, our results revealed voltage gated potassium channel Kv7.2 and cystic fibrosis related CFTR channel modulators significantly inhibit ferroptosis across cell types and inactivated ferroptosis inducers despite the fact that the cell lines did not express those proteins. We observed that these ion channel modulators significantly decreased lipid peroxidation levels across different cell lines and ferroptosis inducers to varying degrees, indicating their role in ferroptosis through independent off-target effects. In contrast, their role in cytoplasmic ion levels was more complex and varied where some modulators surprisingly led to changes in ion levels during the course of ferroptosis through off-target effects. Other inhibitors showed minimal effect on potassium levels, yet all of them could prevent ferroptosis, indicating their inhibitory activity is not directly correlated with the ion levels and their targets leads to downstream effects involved in ferroptosis machinery such as Nrf2 pathway^26^. We also discovered that several known ferroptosis inhibitors such as KRCA008, SHP099, and Seratrodast also influences the ion levels to varying degrees, offering a view where cellular components influence the ion channel activity which in turn affects other cellular functions. Yet, elucidating this exact mechanism would require several studies across multiple disciplines and is beyond the scope of this work which aimed to highlight the overall role of ion homeostasis in ferroptosis.

Because of the delicate ion balance we observed, we hypothesized that the loss of a specific ion type does not lead to enhanced ferroptosis, whereas loss of a specific ion can still rescue a cell from ferroptosis, such as the off-target effect of CFTR inhibitor decreasing potassium levels. Therefore, we hypothesized whether excess changes in cellular ion levels may be compensated through other mechanisms and would activate alternative pathways that prevent cell death. This led us to reason rather than promoting excess ion loss, inhibiting the cellular mechanisms that are essential to ion homeostasis may result in enhanced ferroptosis. This would result in a sustained ion dysregulation that can hinder cell homeostasis. Therefore, we investigated clinically approved Na^+^/K^+^ ATPase inhibitor digoxin in combination with ferroptosis inducers erastin and RSL3. Our approach led to an enhanced cell death in RSL3, but not in erastin when combined with digoxin. Despite these findings, in vivo application of RSL3 is highly limited leading us to test combination of salinomycin and digoxin as an enhanced ferroptosis therapy. Our findings revealed that the combination of digoxin and salinomycin enhances lipid peroxidation and increases cell death across cell lines. This consistent ferroptosis induction and enhanced cell death, encouraged the in vivo investigation of the combination. This combination therapy reduced tumor burden in xenografted mice beyond either therapy alone. Overall, our salinomycin-digoxin combination offers a novel and clinical feasible approach to ferroptosis and highlights the role of ion imbalance as a ferroptosis target. While limited bioavailability of salinomycin required daily injections for observing therapeutic benefits, analogs of salinomycin can offer better clinical outcomes and serve as the future ferroptosis therapeutics. It is important to highlight our results not only offer insights to induce ferroptosis in tumors using clinical drug digoxin, but also identifies two clinically approved agents, ivacaftor^26^ and flupirtine as potent ferroptosis inhibitors. Therefore, our results can lead to drug repurposing and clinical trials utilizing these agents against diseases such as neurodegeneration where ferroptosis is implicated. It is important to highlight digoxin was tested in clinical trials against circulating tumor cells in metastatic breast cancer patients^37^ and previous studies showed the neuroprotective effects of flupirtine Creutzfeldt-Jakob disease^38^. Overall, these studies demonstrate the validity of our findings and highlight where the future work on ferroptosis related therapies can evolve through drug repurposing and utilizing ion homeostasis as a target.

## Methods

### Cell Culture

MDA-MB-435 and HT-1080 cells were obtained from American Type Culture Collection(ATCC). MDA-MB-435 cell line was cultured in RPMI 1640 Medium (Thermo Fisher), 10 % Fetal Bovine Serum (FBS) and 1% Penicilin-Streptomycin (Pen-Strep) Solution. HT-1080 cell line was cultured with DMEM medium (Thermo Fisher) with 10% FBS and 1% Pen-Strep. Media was changed in every 3 days to ensure nutrient availability throughout the cell growth and cells were passaged at 80% confluence with 0.05% Trypsin EDTA (Thermo Fisher). Cell viability and health were constantly monitored during the course of the study to ensure the reproducibility of the assays.

### Cytotoxicity

Cells were trypsinized and seeded on 96-well plate with 10,000 cells per well suspended in 150 μL of their respective media. The cells were incubated in 37 °C with 5% CO_2_ levels for 24 hours for surface attachment. Only the center 60-wells were used for the assays to avoid the edge-effects. The cells were treated with erastin(10 μM) or RSL3(2 μM)(Cayman Chemicals) in the presence and absence of the inhibitors and activators. These modulators were purchased from Cayman Chemicals and MedChem Express. Their source and final concentrations are given in Supplementary Tables. Cells were treated with ferroptosis inducers and modulators for 24 hours and cytotoxicity was measured using Cell Titer Glo2 Assay (Promega) based on the manufacturer’s instructions. The cells were incubated with Cell Titer Glo2 reagent for 10 minutes and this mixture is transferred to a black-walled plate to prevent cross-well light detection. The bioluminescence was measured using Molecular Devices SpectraMax iD5 Multi-Mode Microplate Reader with 1000 ms integration time with open spectral detection.

### Live Cell Calcium Imaging

Cells were trypsinized and seeded on 96-well plate with 10,000 cells per well suspended in 150 μL of their respective media into an imaging plate with glass bottom with cell attachment surface. The cells were incubated in 37 °C with 5% CO_2_ levels for 24 hours for surface attachment. Only the center 60-wells were used to avoid the edge-effects. The cells were treated with Fluo-3 AM (Thermo Fischer) for 30 minutes with 10 μM final concentration and washed with PBS twice and fresh media was added to the wells. The cells were then treated with erastin(10 μM) and imaged using Zeiss Microscope Widefield Axio Observer equipped with incubation chambers that can maintain 37 °C with 5% CO_2_ levels overnight. The cells were imaged every 15 minutes with 488 nm laser excitation over 16 hours to limit phototoxicity. The images were processed using ImageJ software.

### Lipid Peroxidation

Trypsinized cells were placed in 24-well plates with 75,000 cells per well with a volume of 500 μL. The cells were incubated in 37 °C with 5% CO_2_ levels overnight for surface attachment. Each well was treated with erastin (20 μM) for 6 hours or RSL3(2 μM) for 2 hours either with our without the ferroptosis inhibitors. For the last 30 minutes of the incubation, C11-BODIPY^581/591^ (Molecular Probes) (2 μM) dye was added to cell culture media based on manufacturer’s instructions. After the 30-minute incubation with the dye, the cells were washed with PBS and trypsinized for flow cytometry preparation. They were subsequently centrifuged for 5 minutes at 1000 rpm and resuspended in PBS containing 2% FBS. Later, they were centrifuged one more time and resuspended in PBS-2%FBS containing DAPI (1 μM) for 5 minutes and they were analyzed using MACS Quant flow cytometry equipment. We utilized vehicle treated wells as a baseline to measure relative increase in green fluorescence signal which indicates increased lipid peroxidation. The cells were further analyzed using FlowJo program.

### Mitochondrial Superoxide Measurement

Cells were trypsinized and seeded on 24-well plate with 75,000 cells per plate in 500 μL. The cells were incubated in 37 °C with 5% CO_2_ levels overnight for surface attachment. The cells were treated with erastin(20 μM) in the presence and absence of the inhibitors for 6 hours. The cells were then incubated with MitoSOX mitochondrial superoxide indicator dye (4 μM) for 30 minutes and washed with PBS based on manufacturer instructions. The cells were then trypsinized and utilized the same protocol outlined in the lipid peroxidation section above. Cellular red fluorescence signal was measured where an increase indicates higher mitochondrial superoxide levels. These changes in fluorescence were analyzed using the FlowJo program.

### Measuring Cytoplasmic Ion and Membrane Polarization Levels with Flow Cytometry

Cells were trypsinized and were seeded on a 24-well plate with 75,000 cells per plate suspended in 500 μL media. The cells were incubated in 37 °C with 5% CO_2_ levels overnight for surface attachment. Each well was treated with erastin (20 μM) for 6 hours or RSL3 (2 μM) for 2 hours either with or without the ferroptosis inhibitors. The cells were then treated with ion sensitive dyes for 30 minutes. These dyes include IPG-4 AM(Cayman Chemicals) for cytoplasmic potassium(5 μM final concentration), Cell Permeant Sodium Green (Thermo Fischer) for cytoplasmic sodium (5 μM final concentration), MQAE (Thermo Fischer) for cytoplasmic chloride concentration(10 μM final concentration) and DiBAC_4_ (Thermo Fischer) for membrane polarization(5 μM final concentration). They were then washed with PBS and further trypsinized and processed as outlined in the lipid peroxidation section. The samples were then analyzed in the flow cytometer based on manufacturer’s instructions and their changes in fluorescence was monitored. Flowjo program was utilized to quantify these changes and gates were designed to report differential changes of fluorescence such as decrease in IPG-4 AM signal with loss of potassium or increase of MQAE signal with the loss of chloride ions as MQAE is quenched by chloride. The results were analyzed using GraphPad Prism software where we calculated the means and standard deviations of each group and performed one-way Anova with multiple comparisons or t-tests where applicable as detailed in each figure legend.

### In vivo Xenograft Mouse Therapy Experiments

All mouse handling, experimentation, imaging, and housing was performed according to NIH guidelines and via approved IACUC protocols at MSKCC. Tumor xenografts were established with a subcutaneous inoculation of 5.0x10^6^ HT-1080 cells (100 µL of a 1:1 Matrigel-PBS mixture) into the flank of female FoxN1^nu^ mice (6-8 weeks). The mice were randomized to different treatment groups. Tumor growth was monitored with caliper measurements and mouse weight monitored for toxicity. Digoxin (0.5 mg/kg) was given i.p. biweekly, and Salinomycin (2.5 mg/kg) given i.p. daily. Mice were euthanized when tumors exceeded 1 cm3 or mice loss > 20% of their body weight.

## Data Availability

All data supporting the findings of this work are available within the article and its Supplementary Information.

## Acknowledgments

We would like to thank RARC at MSKCC for their guidance in animal experimentation and husbandry, as well as the MSKCC Antitumor Assessment core, particularly Sydney Bowker, for assistance with in vivo therapy administration. Support for this work was possible thanks to the NCI R01-CA-257811 and NIBIB R56 EB030512, NCI Cancer Center core grant P30 CA08748 (to Selwyn Vickers), along with Vince Lombardi Cancer Foundation.

## Author Contributions

A. Y. S. designed the experiments with input from E. A. . A.Y. S. and E. A., performed the in vitro experiments and data analysis. A.Y. S., E. A. and N.M. performed the in vivo experiments. A.Y. S., E. A. and N.M. analyzed the data and created the figures 1-3. J. G. supervised the overall work. All authors contributed to the writing and revision of the manuscript.

