## Supplemental data for "Sustained Ion Imbalances Enhance Ferroptosis"

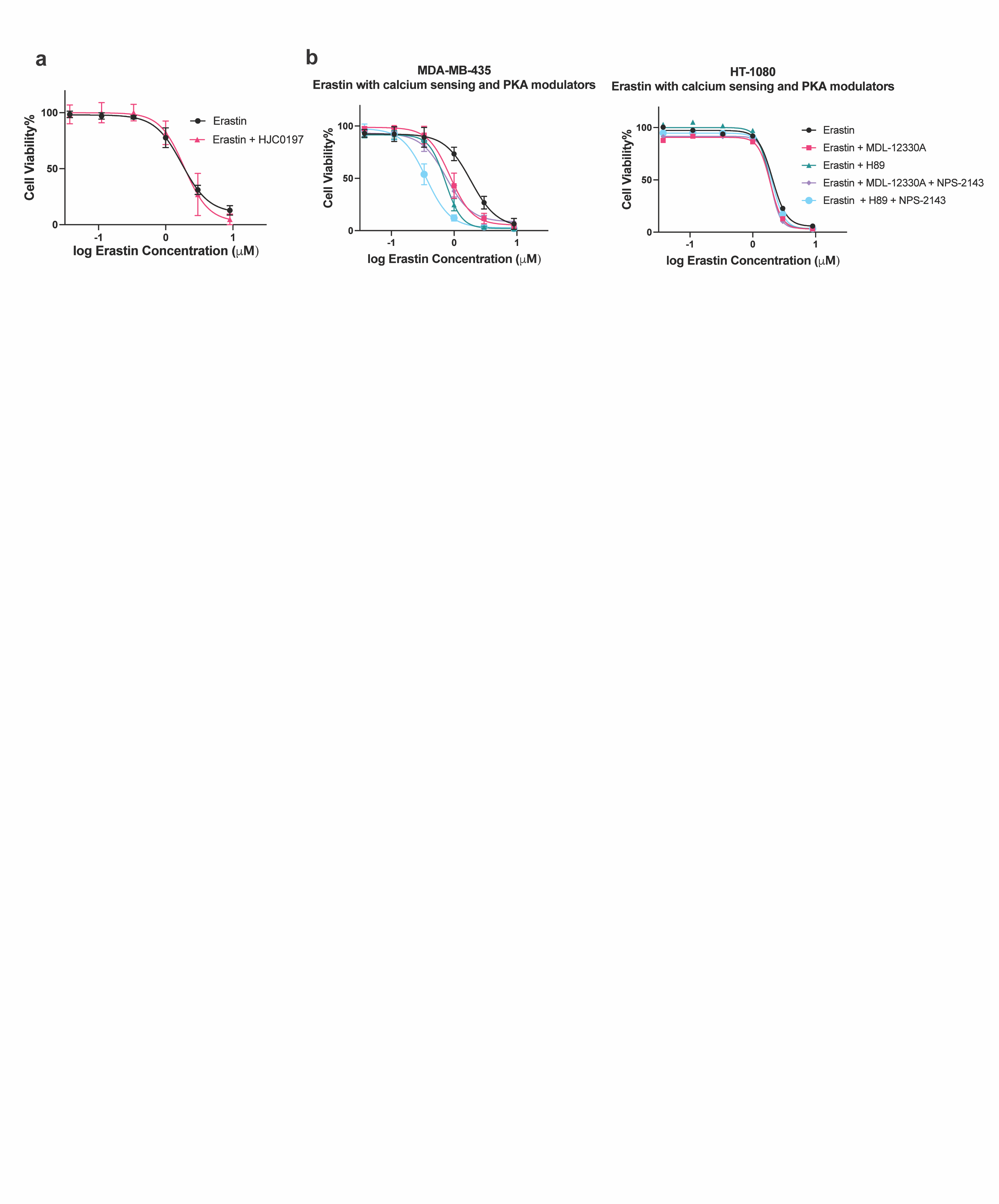

**Supplementary Figure 1. Combining Calcium sensing and PKA modulators leads to enhanced cell death in specific cell lines a)** EPAC inhibitor HJC0197 (5 μM)	has no effect on erastin induced cytotoxicity,(Erastin IC50=1.742 μM, Erastin-HJC0197 IC50=1.887 μM**) b)** Combination of Calcium sensing Receptor inhibitor with PKA and Adenylyl Cyclase Inhibitors (5 μM each)	leads to enhancement in erastin induced cell death in MDA-MB-435 cells (Erastin IC50= 1.843 μM, Erastin-MDL-12330A IC50=0.8623 μM, Erastin-H89 IC50=0.7262 μM, Erastin-MDL-12330A-NPS-2143 IC50=0.7499 μM, Erastin H89-NPS-2143 IC50=0.3603 μM)**,** but not in HT-1080 cell line (Erastin IC50=2.051 μM, Erastin-MDL-12330A IC50=1.905 μM, Erastin-H89 IC50=1.834 μM, Erastin-MDL-12330A-NPS-2143 IC50=1.912 μM, Erastin H89-NPS-2143 IC50=2.020 μM). For cytotoxicity assays n=3 biological replicates, Mean ± s.d. In kill curves concentrations were converted to the log scale and the values were fitted as a nonlinear fit using the log(inhibitor) vs. response function with variable slopes and four parameters.

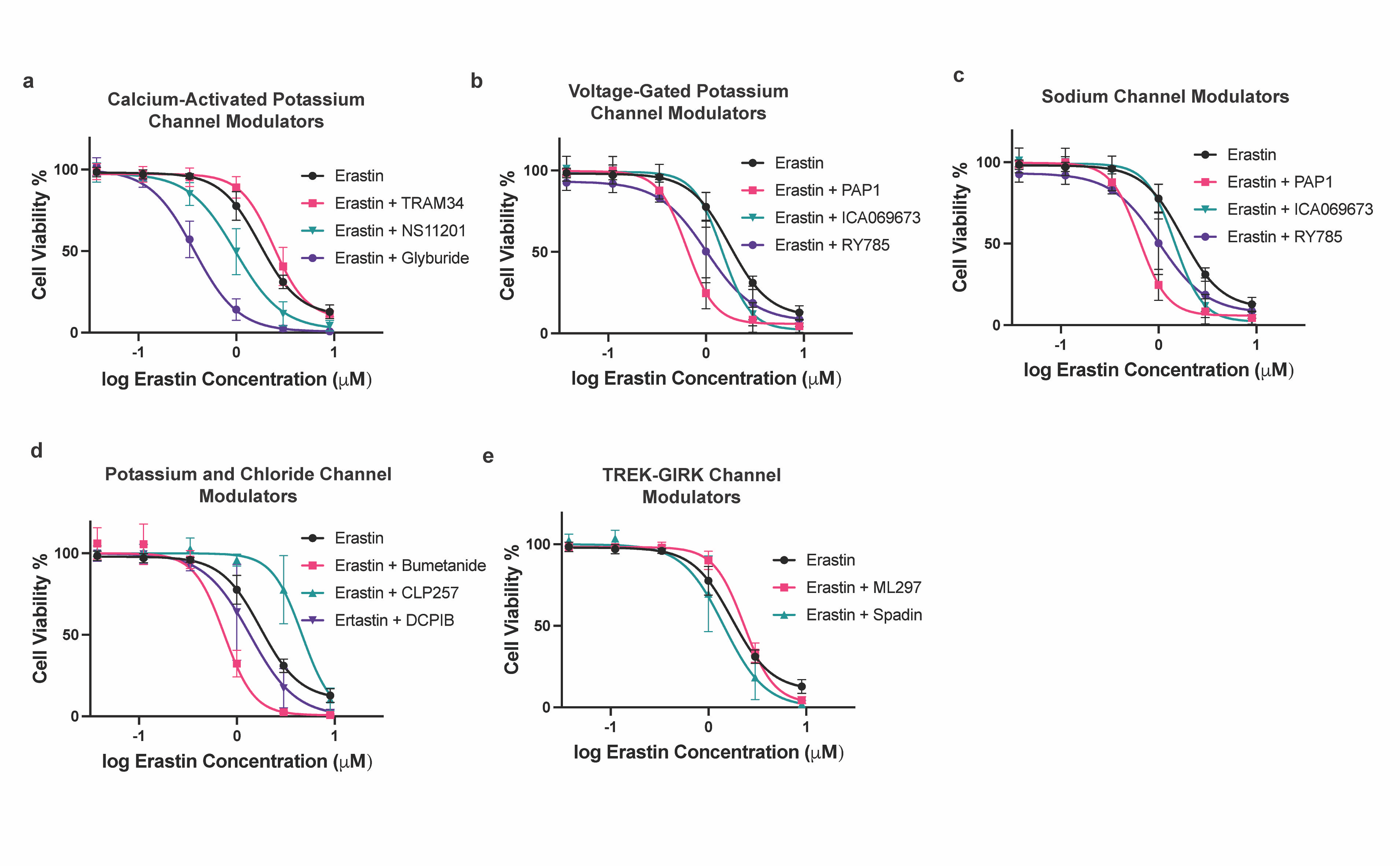

**Supplementary Figure 2. Role of different ion channel modulators on erastin induced ferroptosis demonstrates the importance of maintaining ion balance for cell survival** Combination of erastin with ion channel modulators leads to diverse effects depending on the ion channel identity **a)** Calcium Activated Potassium Channel Modulators, Erastin IC50= 1.742 μM, TRAM-34 IC50= 2.402 μM, NS11201 IC50= 0.9802 μM, Glyburide IC50= 0.3712 μM**, b)** Voltage gated Potassium Channel Modulators, PAP-1 IC50= 0.6327 μM, ICA069673 IC50= 1.460 μM, RY785 IC50= 1.010 μM, **c)** Sodium Channel Modulators, Amiloride IC50= 1.431 μM , Ivabradine IC50= 0.9135 μM , CGP37157 IC50= 1.111 μM , **d)** Potassium and Chloride Channel Modulators, Bumetanide IC50= 0.7416 μM , CLP257 IC50= 4.638 μM , DCPIB IC50= 1.363 μM, **e)** TREK-GIRK ion channel modulators ML297 IC50= 2.315 μM , Spadin IC50= 1.435 μM. For cytotoxicity assays n=3 biological replicates, Mean ± s.d. In kill curves concentrations were converted to the log scale and the values were fitted as a nonlinear fit using the log(inhibitor) vs. response function with variable slopes and four parameters.

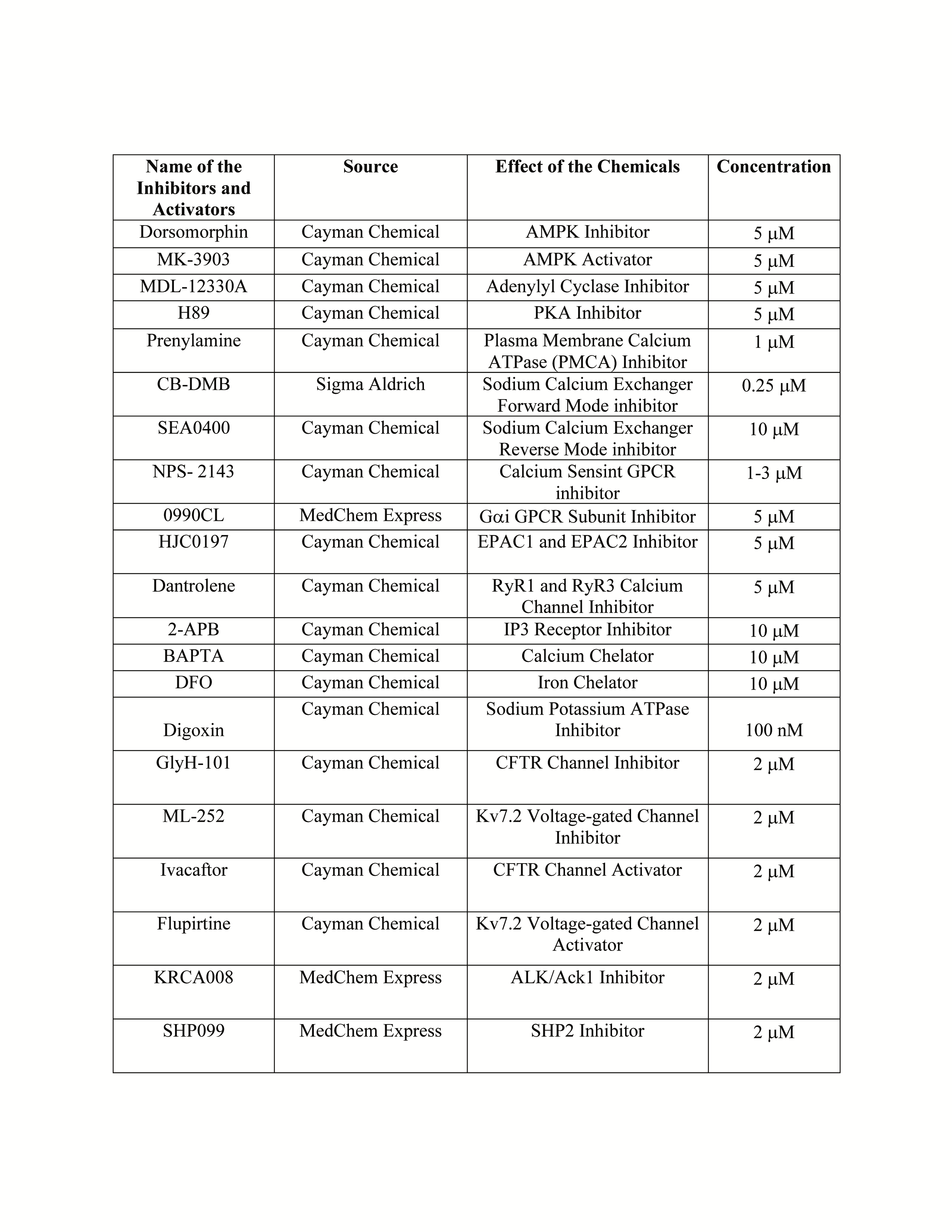

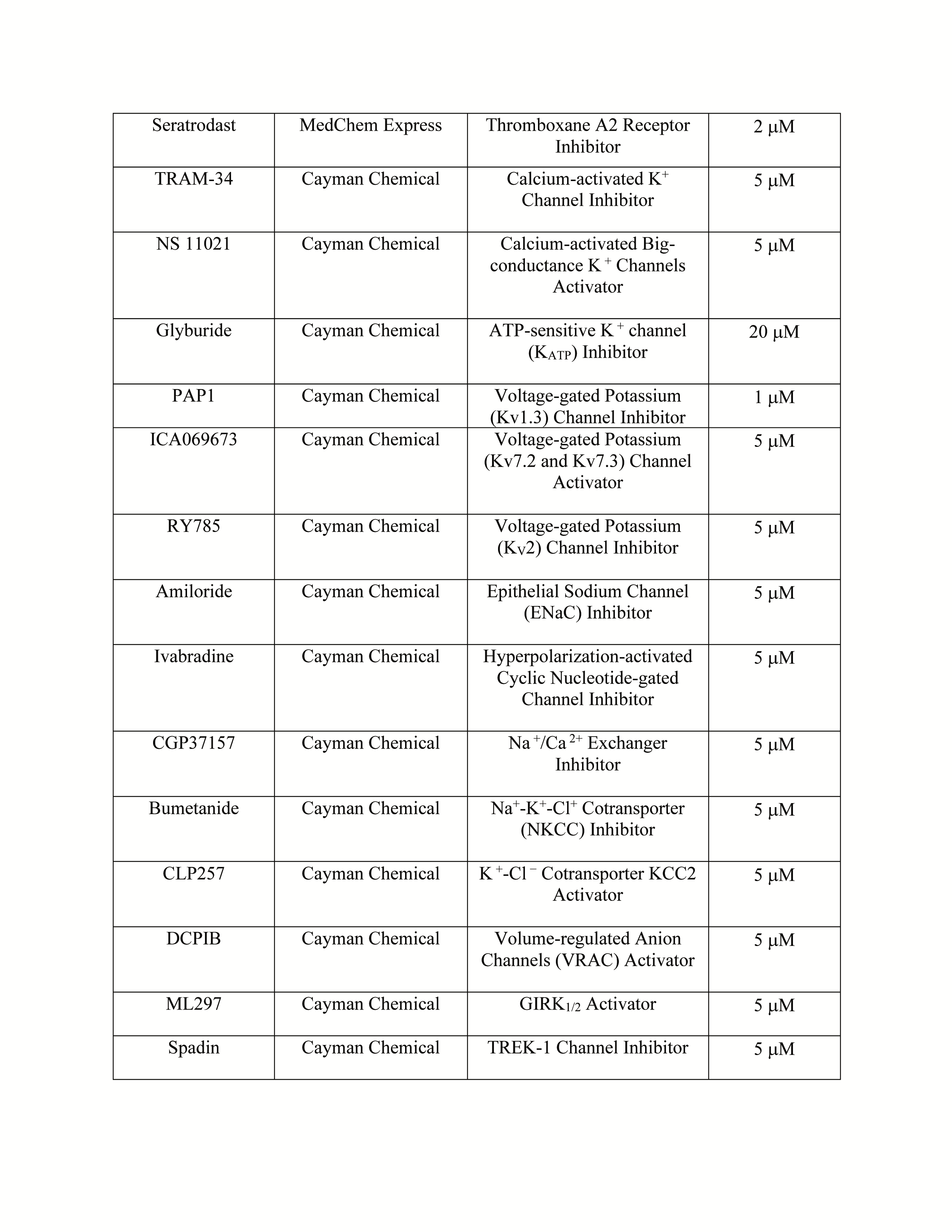

**Supplementary Figure 3. Table showing the mechanism of action and concentration of different modulators used in combination with erastin** Mechanism of Action for different modulators used in this study along with their concentration.

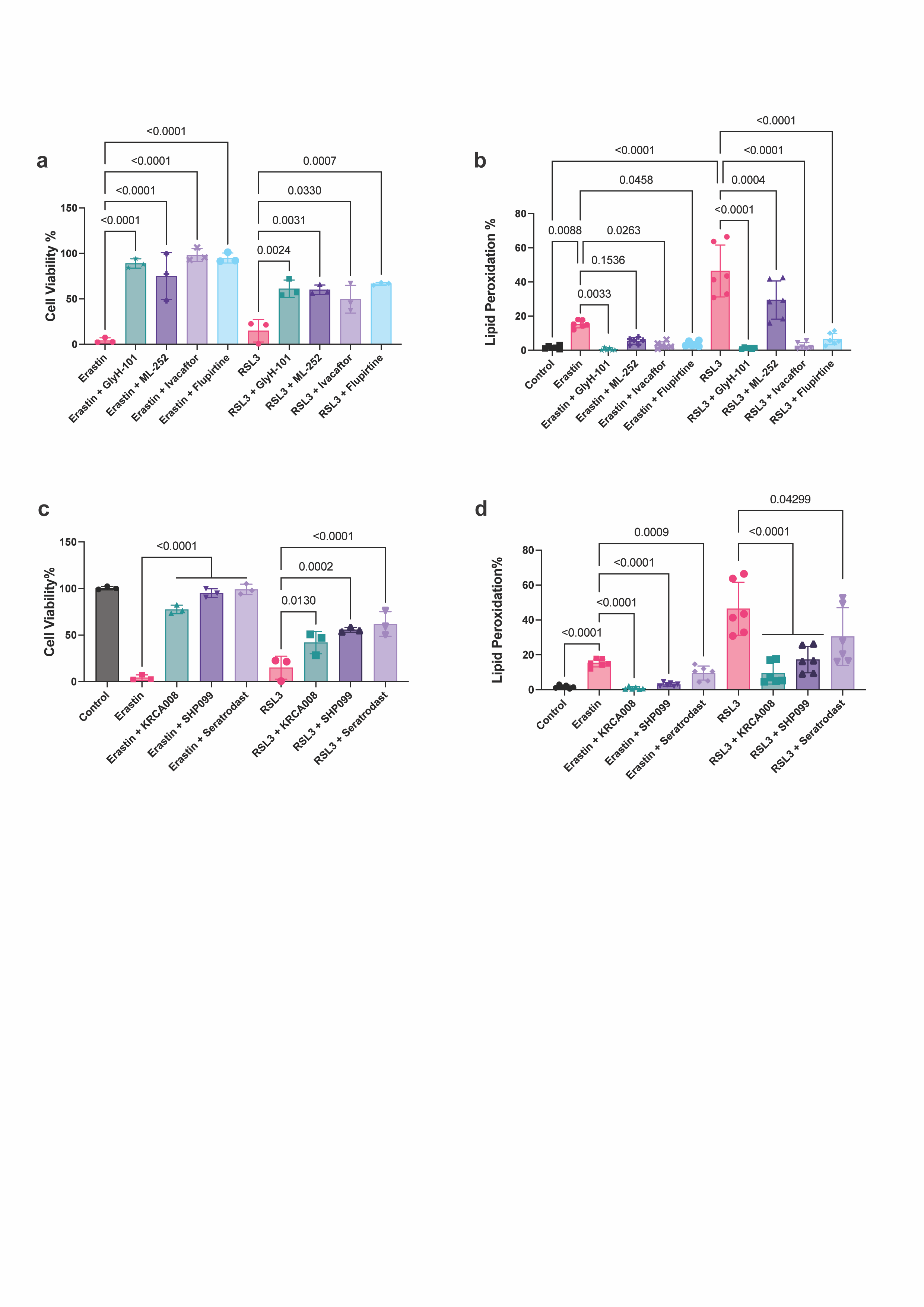

**Supplementary Figure 4. Modulators of CFTR and Kv7.2 and kinase inhibitors can inhibit both erastin and RSL3 induced cell death in HT-1080 cell lines and lead to decreased lipid peroxidation a)** Cytotoxicity of HT-1080 cells with erastin and RSL3 induced cell death is inhibited by GlyH-101 **(2 μM)**	and activator Ivacaftor**(2 μM)**	, as well as Kv7.2 inhibitor ML252 **(2 μM)**	and activator Flupirtine **(2 μM)** **b)** CFTR and Kv7.2 modulators leads to decreases in the lipid peroxidation levels , **c)** Erastin and RSL3 induced cell deaths are inhibited with ALK/Ack1 inhibitor KRCA008 **(2 μM)** , SHP2 inhibitor SHP099 **(2 μM)**	, Thromboxane A2 receptor Seratrodast **(2 μM)**	**d)** Erastin and RSL3 induced lipid peroxidation are modulated by KRCA008, SHP099, Seratrodast**,** For cytotoxicity assays n=3 biological replicates, for the lipid peroxidation, potassium and chloride measurements, N=6 pooled from 3 independent experiments. Mean ± s.d. p-values are reported above the lines (Ordinary one-way ANOVA with Tukey’s multiple comparisons).

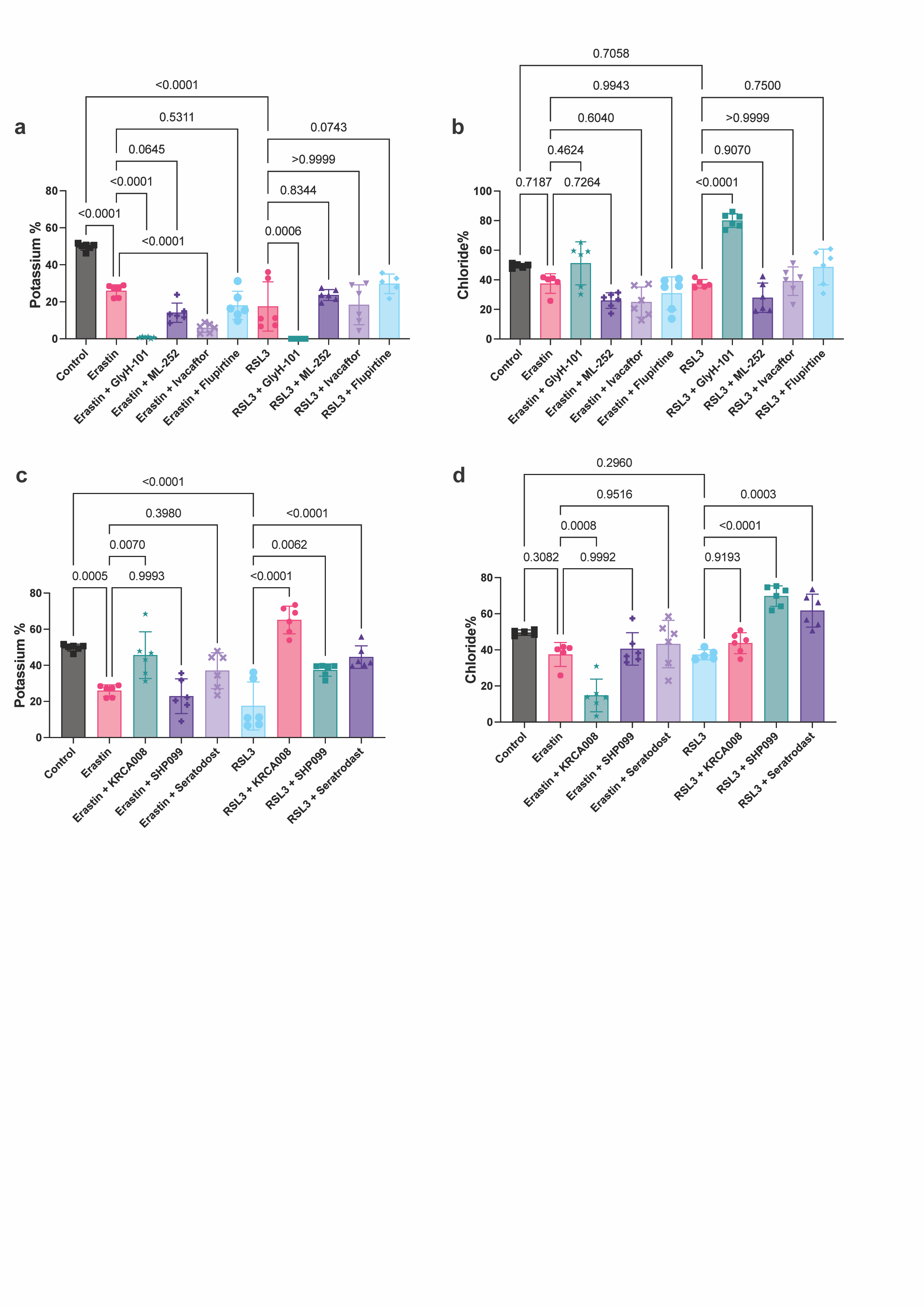

**Supplementary Figure 5. Ionic imbalances in ferroptosis induction with erastin and RSL3 and the effect of CFTR and Kv7.2 modulators and kinase inhibitors in HT-1080 a)** CFTR and Kv7.2 modulators leads to changes in the potassium levels, **b)** CFTR and Kv7.2 modulators leads to changes in the chloride levels in RSL3 induced cell death, but not erastin induced cell death, **c)** Erastin and RSL3 induced potassium level changes are modulated by KRCA008, SHP099, Seratrodast**. d)** Erastin and RSL3 induced chloride level changes are modulated by KRCA008, SHP099, Seratrodast**.** For measuring potassium and chloride levels, N=6 pooled from 3 independent experiments. Mean ± s.d. p-values are reported above the lines (Ordinary one-way ANOVA with Tukey’s multiple comparisons).

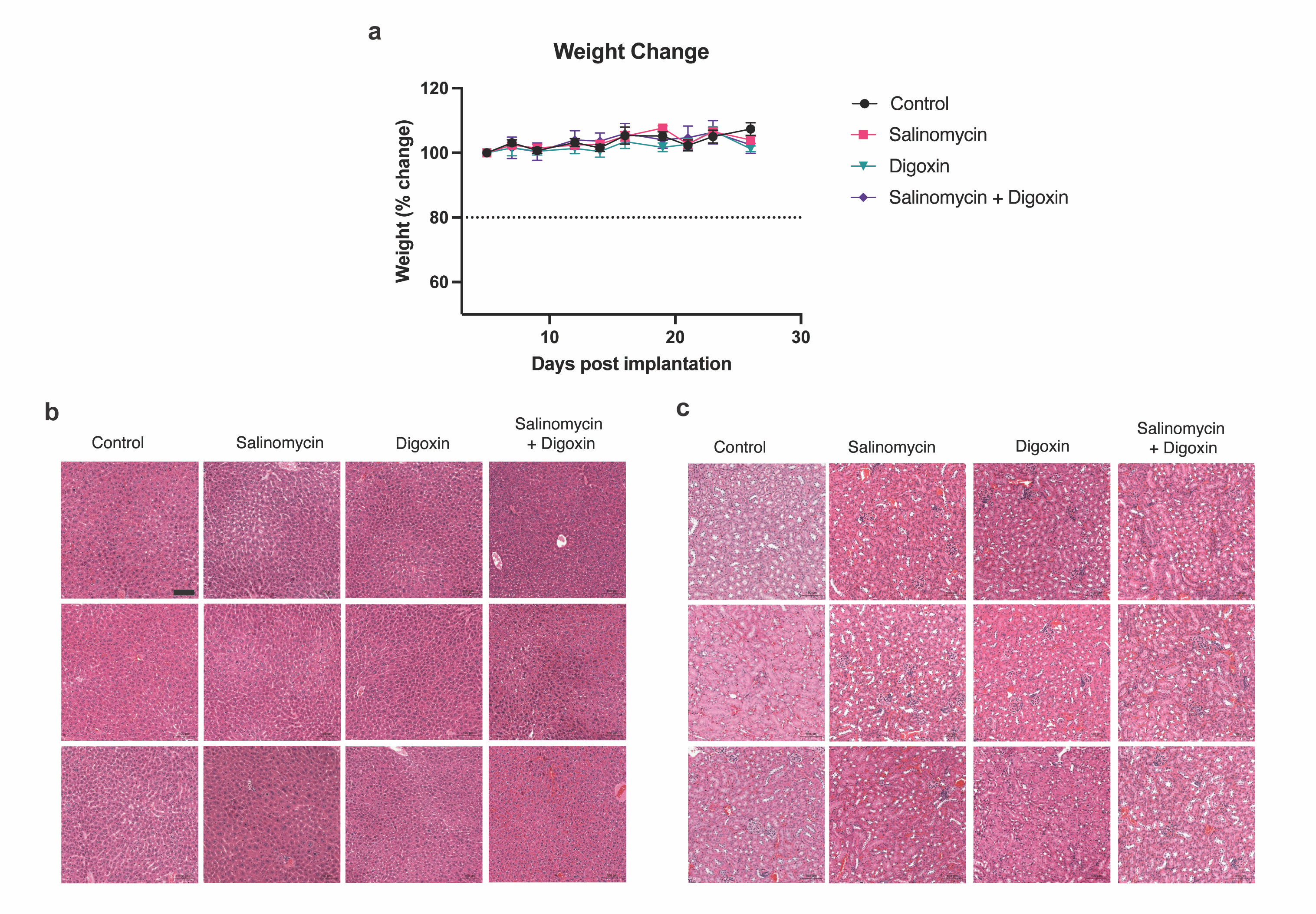

**Supplementary Figure 6. Salinomycin-Digoxin combination does not influence the overall health of mice. a)** Salinomycin alone, Digoxin alone or their combination therapy does not influence the mouse weight during the course of the therapy **b)** Salinomycin alone, Digoxin alone or their combination therapy does not harm the liver tissue based on H&E staining **c)** Salinomycin alone, Digoxin alone or their combination therapy does not harm the kidney tissue based on H&E staining, N=10 mice for each group.
